# BARseq3: a modular system for integrating spatial multi-omics and cellular barcoding in single cells

**DOI:** 10.64898/2026.05.13.724900

**Authors:** Huihui Qi, Manjari M-G Anant, Dylan Z. Faltine-Gonzalez, Ruitao Hu, Lai Wei, Christopher D. Workman, Caleb Shi, Ishbel Del Rosario, Justus M. Kebschull

## Abstract

Understanding cellular identity requires multimodal measurements in single cells. Cellular barcoding provides powerful tools for recording the properties or history of individual cells in nucleic acids, while spatial omics techniques enable the measurement of a growing list of molecular features at micron resolution in tissue. However, existing methods that integrate these approaches in single samples are limited in the modalities they support, their flexibility, and efficiency. Here, we present BARseq3, a modular system that combines cellular barcoding with high-efficiency spatial transcriptomics and translatomics at subcellular resolution in tissue. BARseq3 is compatible with fixed samples, immunostaining, diverse species, and can be easily extended to include other spatial assays, enabling a multimodal understanding of cellular identity.

## MAIN

Cellular identity is commonly defined by the transcriptomic state of a cell: the set of mRNAs present at the moment of measurement.^1–4^ However, this transcriptomic definition captures only one dimension of the cellular phenotype. Other features also influence cellular identity, including location, post-transcriptional regulation of mRNAs, developmental lineage, stimulus response history, and, in the case of neurons, connectivity.^5^ As these different features are often partially or completely orthogonal to each other,^6,7^ a comprehensive understanding of cellular identity must rely on the intersection of distinct cellular features.

Different cellular features are traditionally measured in separate experiments and tissue samples for technical reasons. Recently, advances in spatial omics methods have enabled readout of gene expression or translation across 3D space in tissue sections.^8–13^ Cellular barcoding approaches, in which individual cells are tagged with unique nucleic acid sequences, now enable the recording of additional, orthogonal cellular features in RNA space, including lineage, neuronal connectivity, or cellular histories of stimuli of interest.^14^ The combination of cellular barcoding and spatial omics in single cells thus promises a truly multimodal readout of cellular identity. However, this integration is complicated by the structure of the barcode. In many methods, the barcode sequence is unknown, short, and/or variable, such as random 30-mers^15,16^ or CRISPR scars.^14,17,18^ These barcodes are unsuitable for identification by the hybridization-based spatial omics methods currently dominant in the field.

To overcome this bottleneck, pioneering padlock-based methods,^19^ Baristaseq,^20^ and BARseq^6^ sequence unknown barcodes directly in tissue sections without assumptions on sequence identity, and BARseq2 integrates this *de novo* barcode sequencing with padlock-probe-based spatial transcriptomics.^21^ *De novo* sequenced barcodes can then be combined with barcoded connectomics methods, such as MAPseq,^15,22^ to link transcriptomic cellular identity with neuronal connectivity at single neuron resolution.^21^ This approach has already begun to shed light on how these two modalities interact in the mouse brain to define integrated connectomic-transcriptomic neuronal cell types.^6,21^ Similarly, integrating lineage barcodes and transcriptomics would reveal how lineage history shapes acute transcriptomic identities. However, in these previous methods, barcode and gene readout are experimentally tightly coupled. As a result, integrating the barcode readout with modalities beyond transcriptomics is challenging, preventing the assessment of how the barcoded modality corresponds to other molecular measures. At the same time, gene detection efficiency within these methods is relatively low, limiting the resolution with which transcriptomic variation can be measured.^21^ To address this, we developed BARseq3, which decouples barcode sequencing from the spatial detection of other modalities and enables plug-and-play coupling of barcoding to high-efficiency, hybridization-based spatial omics, including *in situ* transcriptomics and translatomics, in the same cells.

BARseq3 is implemented as a modular, three-part workflow that is applied to tissue slices that contain cellular barcodes introduced, via, for example, viral or genomic delivery, depending on the cellular barcoding assay (**Fig. 1**). In the Gene Module (**Fig. 1A**), gene expression information such as the transcriptome and/or translatome is captured in dedicated libraries directly on endogenous mRNA by hybridizing oligonucleotide ^NH^ probes (for example, SNAIL or TRI probes adapted from STARmap^10^ and RIBOmap^8^). Bound probes are then circularized, amplified by rolling circle amplification (RCA), and covalently tethered to the tissue. In the Barcode Module (**Fig. 1B**), adapted from BARseq,^6^ barcode information is captured into a another, separate library. Barcode RNA is reverse transcribed and probed with a gapped padlock. The gap is filled, and the padlock is circularized, amplified by RCA, and tethered to the tissue. Finally, in the Sequencing Module (**Fig. 1C**), gene IDs in gene amplicons and barcode sequences in barcode amplicons are determined *in situ* by Illumina sequencing-by-synthesis chemistry.^20^ Each Gene and Barcode Module library is sequenced independently and sequentially using orthogonal sequencing primers. Illumina sequencing chemistry allows for efficient long-read (>30 nt) sequencing and more standardized, robust reagents than sequencing-by-ligation chemistries typically employed in *in situ* sequencing methods.^8,10^ Importantly, in BARseq3 the Gene and Barcode Modules and their associated Sequencing Module are independent of each other. As a result, BARseq3 is flexible to accommodate alternative or additional RCA-based spatial omics methods in the Gene Module without redesigning the barcode capture chemistry and vice versa. Similarly, different Gene Modules can be run with or without the Barcode Module, and the Barcode Module without a Gene Module, depending on what cellular feature should be measured in an experiment (**Fig. 1D**).

**Fig. 1:**
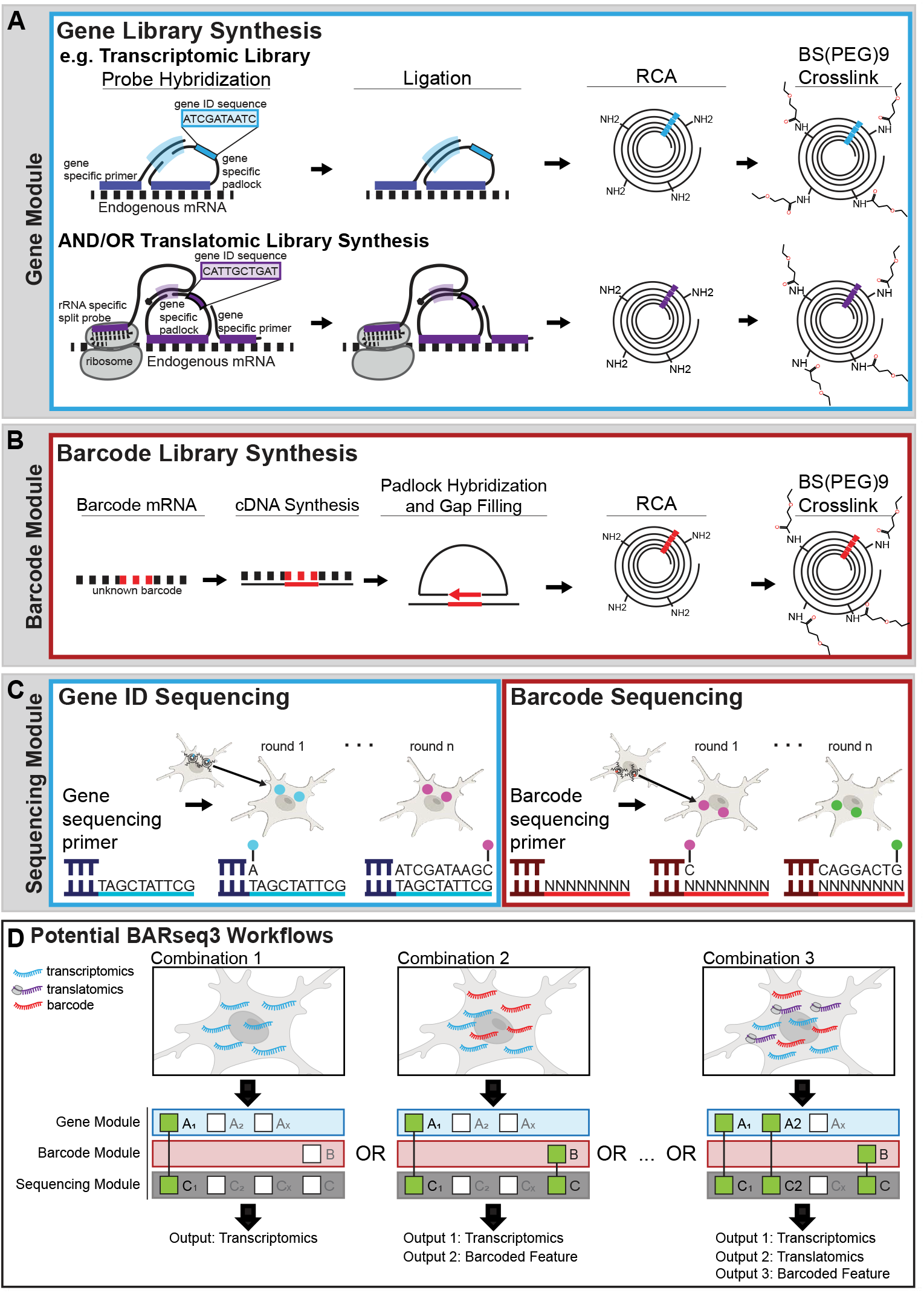
The modular BARseq3 workflow. **(A)** The Gene Module can incorporate a variety of hybridization-based spatial omics assays. A transcriptomic (top) and a translatomic (bottom) library are shown. Assay-specific probes, including a padlock probe containing a gene ID sequence, are hybridized to endogenous RNA, circularized, amplified, and cross-linked to the tissue. **(B)** The Barcode Module relies on a gapped padlock probe that hybridizes to known sequences flanking the unknown barcode on cDNA. The gap is then filled and the padlock is ligated, amplified, and cross-linked. **(C)** The Sequencing Module utilizes Illumina sequencing-by-synthesis chemistry. Library-specific sequencing primers allow independent sequencing of gene IDs in different Gene Module libraries and unknown barcodes in Barcode Module libraries. **(D)** BARseq3 is highly modular. Experiments can utilize the Gene Module only, one gene modality or multiple, and in combination with the Barcode Module, yielding single experiments that report many kinds of information.

To assess BARseq3 performance, we first tested the integration of transcriptome and barcode detection. We delivered random 30-nucleotide-long cellular barcodes to neurons in mouse motor cortex using a library of barcoded Sindbis viruses.^23^ Barcodes of this type are used in the barcoded connectomics method MAPseq,^15^ which relies on cellular barcoding to read out neuronal projections. We then performed BARseq3, probing for genes *Slc17a7* and *Gad1* with four STARmap-style SNAIL probes^10^ per gene and the unknown cellular barcodes. In parallel, we subjected adjacent brain sections to the BARseq2 protocol, probing the same genes with 12 padlocks per gene as well as the unknown barcodes (**Fig. 2A**). BARseq3 yielded bright gene and barcode amplicons, with barcode signal filling infected cells as expected for the highly expressed Sindbis barcodes (**Fig. 2B**). Illumina sequencing of BARseq3 gene amplicons showed good signal-to-noise (**Sup Fig. 1A, B**) and low lag (**Sup Fig. 1E,F**). Similarly, BARseq3 barcode sequencing signal-to-noise was high across 10 rounds of sequencing (**Fig. 2C,D**). BARseq3 is also compatible with PFA-fixed tissue after Pepsin pretreatment (**Sup Fig. 2**).

**Fig. 2:**
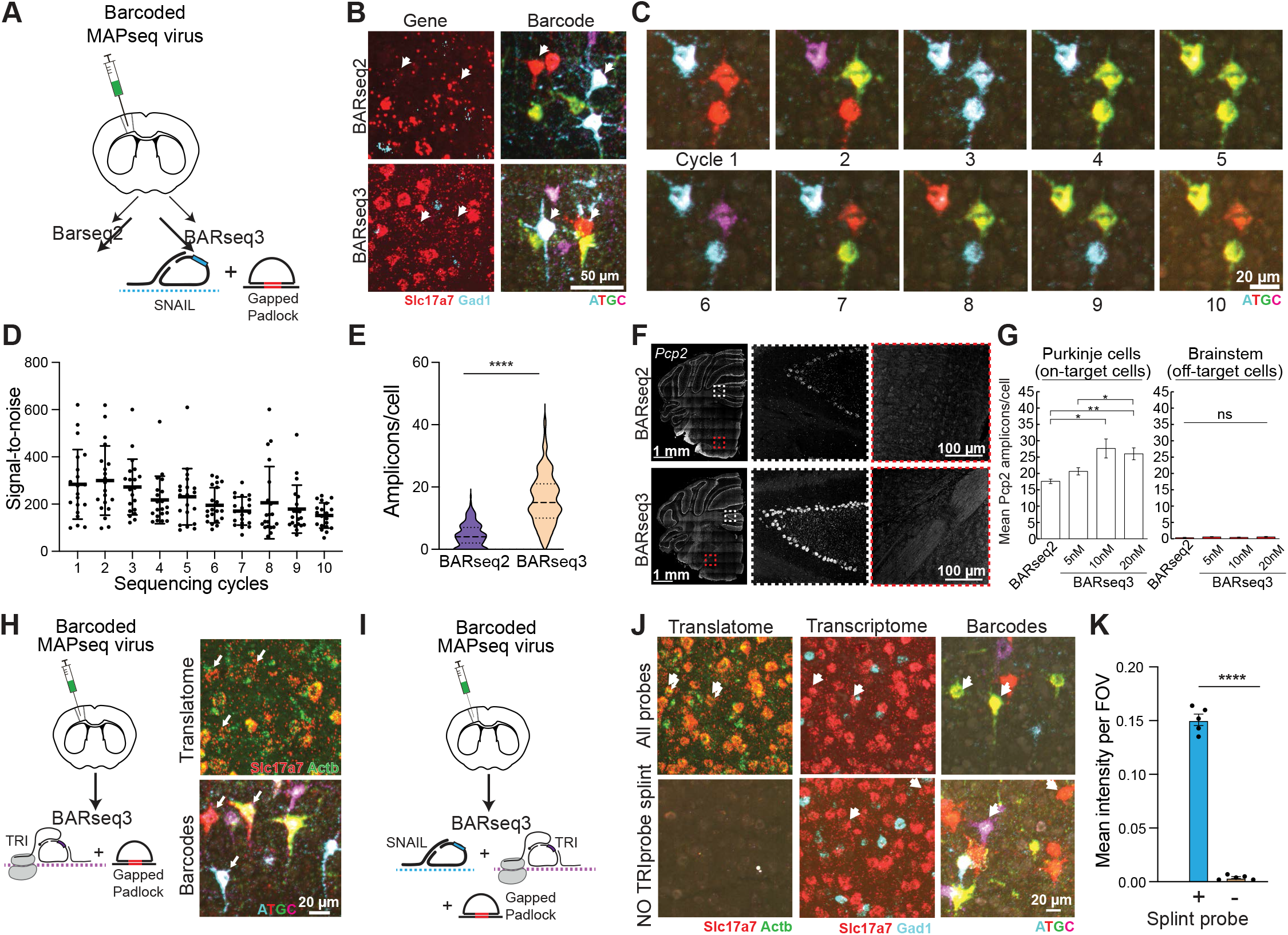
BARseq3 improves gene-detection sensitivity and enables plug-and-play multi-omics with cellular barcoding. **(A)** Schematic of the experimental design for comparing BARseq2 and BARseq3. **(B)** Representative gene (*Slc17a7* and *Gad1*) and barcode sequencing images for the two methods. **(C)** Representative images of 10 sequential BARseq3 barcode sequencing cycles. **(D)** Signal to noise of BARseq3 barcodes remains high across ten sequencing cycles. **(E)** BARseq3 produces more amplicons than BARseq2 in barcoded cells (BARseq2 cells n=229, BARseq3 cells n=360, Student’s t-test, ****pvalue < 0.0001). (**F)** Representative images of the *Pcp2* specificity experiment (10 nM), highlighting the on-target Purkinje cell signal and the off-target brainstem signal for each method. **(G)** *Pcp2* on-target signal is higher for BARseq3 than BARseq2, and increases with BARseq3 probe concentration (n=4, FOVs each, Welch’s t-test with Bonferroni correction, *pvalue < 0.0083, **pvalue < 0.0016). Off-target amplicons are rare and not significantly different across methods or probe concentrations concentration (n=4, FOVs each, Welch’s t-test with Bonferroni correction, *pvalue < 0.0083). (**H)** Experimental design for simultaneously probing translatome and barcodes (left) and representative images showing BARseq3 translatome library amplicons within barcoded cells (right). **(I)** Experimental design for simultaneously probing the translatome, transcriptome, and barcodes. **(J)** Representative images highlighting transcriptomic and translatomic amplicons within barcoded cells. No translatomic amplicons are formed when the ribosome-bound splint probe is omitted. **(K)** Quantification of the *Slc17a7* translatome amplicons signal intensity with and without the splint probe (n=5 FOVs each, Student’s t-test, ****pvalue < 0.0001).

When we compared BARseq2 and BARseq3 gene signals, we found that BARseq3 produced more gene amplicons per cell than BARseq2 in both barcoded (**Fig. 2E**) and uninfected (**Sup. Fig. 1F**) neurons despite using 67% fewer probes per gene and lower concentrations per probe, indicating improved sensitivity. To measure BARseq3 gene signal specificity, we first validated that *Slc17a7* and *Gad1* gene amplicons were confined to different cells, as expected (**Sup. Fig. 1G**). To more directly assess specificity, we then took advantage of the highly localized gene expression of *Pcp2* to cerebellar Purkinje cells in the hindbrain (**Fig. 2F**) and *Malat1* to nuclei (**Sup. Fig. 3**) and measured signal from these genes in ‘on-target’ (Purkinje cells/nuclei) and adjacent ‘off-target’ (brainstem tissue/non-nuclei) regions in parallel BARseq2 and BARseq3 experiments. We found that on-target BARseq3 signal for *Pcp2* was probe concentration dependent and higher than that of BARseq2 (**Fig. 2G**), while off-target signal was low and not significantly different between methods and conditions. Similarly, for *Malat1* and at all concentrations tested, the ratio of on-target to off-target signal was probe concentration dependent and significantly higher in BARseq3 than BARseq2 (**Sup. Fig. 3B**). Taken together these results demonstrate improved sensitivity and specificity of BARseq3 over BARseq2 in combining barcode and transcriptome readout in single cells, but also highlight the need to use an appropriate probe concentration for optimal signal-to-noise.

**Fig. 3:**
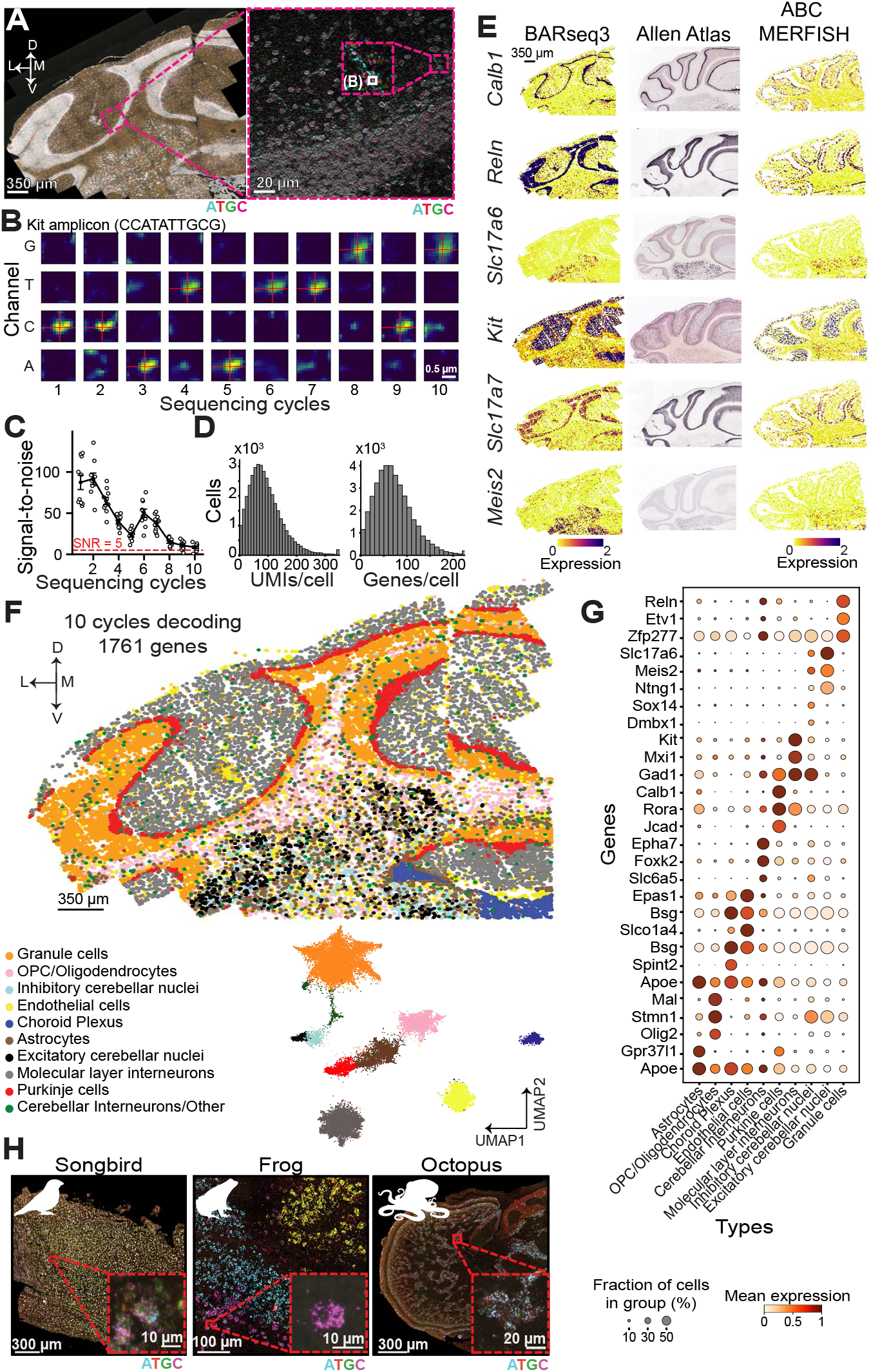
High-resolution BARseq3 transcriptomic assay in the adult mouse cerebellum. **(A)** BARseq3 imaging of sequencing cycle 1 in an adult mouse cerebellar coronal section. Left: low-magnification overview of all fields of view. Right: One field of view showing individual amplicons/cells. **(B)** Example amplicon assigned to *Kit*. Four-channel base imaging across 10 sequencing cycles shows the expected cycle-by-cycle color changes for the called gene ID. **(C)** Amplicon signal-to-noise ratio is high across 10 sequencing cycles and remains in the decodable range of more than 5 standard deviations across background, as defined by the Rose criterion. **(D)** Histograms of number of UMIs per cell (mean: ~92) and number of unique genes per cell (mean: ~71). **(E)** Marker gene expression patterns compared across technologies: BARseq3, Allen ISH Atlas, and Allen Brain Cell MERFISH Atlas. BARseq3 shows higher signal while preserving expected spatial patterns. **(F)** Spatial map of inferred cerebellar cell types (10 cycles; 1,745 genes), showing canonical laminar/region-specific localization (top) and corresponding UMAP embedding colored by type (bottom). **(G)** Dot plot of marker gene expression across cell types, consistent with known cerebellar markers. **(H)** Representative images of BARseq3 transcriptomic signal (sequencing cycle 1) in songbird, frog, and octopus brain tissues.

To demonstrate the modularity of BARseq3, we next swapped the SNAIL probes for RIBOmap-style TRI probes^8^ to measure a different cellular modality. These probes produce amplicons only on actively translated mRNAs (**Fig. 2H**). We probed barcoded mouse motor cortex with TRI probes against *Slc17a7* and *Actb* and padlocks against the unknown barcode and performed BARseq3. Again, we observed good signal-to-noise gene amplicons and strong barcode signals (**Fig. 2H**; **Sup Fig 1B**). To test our ability to measure multiple cellular features in single cells within the Gene Module, we then probed barcoded tissue sections with both SNAIL and TRI probes as well as barcode probes and performed BARseq3 (**Fig. 2I**). This experiment yielded strong signal for total mRNA from SNAIL probes, translating mRNA from TRI probes, and barcodes from the gapped padlock probes in the same tissue section (**Fig. 2J)**. Importantly, no translatome amplicons were formed when the ribosome-bound splint TRI probe was omitted, validating that the translatome assay specifically detects actively translated mRNA (**Fig. 2K**). Taken together, these data demonstrate that BARseq3 is a flexible system that integrates high-efficiency spatial transcriptomics and translatomics with *de novo* barcode sequencing at subcellular resolution in single cells.

The modular design of BARseq3 allows the use of the Gene and Sequencing Modules without the Barcode Module, yielding, for example, an efficient spatial transcriptomics assay with a robust Illumina sequencing readout (**Fig. 1D**). To validate such an application, we designed a panel of SNAIL probes against 1,745 genes, including all known transcription factors, genes involved in cellular migration and adhesion, and a selection of neuronal marker genes. We probed sections of non-barcoded adult mouse cerebellum at an optimized probe concentration and amplification time, then sequenced 10 cycles using high-resolution imaging (**Fig. 3A,B**). Signal was high across all sequencing cycles, and individual amplicons changed color in patterns consistent with the designed gene ID sequences (**Fig. 3B,C**). We decoded the gene identities of amplicons and assigned them to cells using a custom pipeline based on Bardensr.^24^ On average, we decoded 92 amplicons and 71 genes per cell (**Fig. 3D**), which is on par with comparable spatial transcriptomics assays.^25–27^ Expression patterns of individual genes aligned with the Allen In Situ Hybridization Atlas^28^ and ABC atlas MERFISH datasets^1^ (**Fig. 3E**). Clustering of BARseq3 gene expression data further revealed the expected cerebellar cell types (**Fig. 3F,G**). Reproducibility of gene expression across adjacent serial sections was high (Pearson r = 0.98; **Sup Fig. 4**), and the protocol allows for post-hoc immunolabeling (**Sup Fig. 5**). Overall, BARseq3 achieves accurate, reproducible, and sensitive spatial gene decoding across many genes and diverse cell types.

Finally, to test the versatility and robustness of BARseq3, we used it to detect gene expression in a broad range of species. Modifying only SNAIL probe sequences to match each organism’s transcriptome, our protocol yielded bright gene amplicons in zebra finch (*Taeniopygia guttata*), frog (*Xenopus tropicalis*), and octopus (*Octopus bimaculoides)* brains, showing that the core chemistry of BARseq3 works across the animal kingdom (**Fig. 3H**).

In summary, BARseq3 experimentally decouples gene readout from barcode detection and thus provides a modular system in which different spatial omics technologies can be combined with cellular barcoding to yield multimodal measures of cellular identity. We demonstrate simultaneous, high-efficiency readout of four cellular features — transcriptomics, translatomics, viral barcodes reporting (e.g., neuronal connectivity), and cellular location — in single cells from a variety of animals, with direct applications in brain mapping. Looking ahead, we expect the Gene Module to be readily extendable to additional hybridization-based assays, such as time-resolved RNA labeling approaches (e.g., TEMPOmap^29^). Similarly, the Barcode Module is applicable beyond viral barcodes to genomic lineage barcodes in developmental tracing and to other sequence tags, such as Cas9 sgRNAs in pooled perturbation screens.^30,31^ BARseq3 thus provides an extensible method for spatially integrating disparate modalities at the single-cell level, enabling a truly multimodal understanding of cellular identity.

### ONLINE METHODS

#### Animals

We conducted all procedures in accordance with the Johns Hopkins University Animal Care and Use Committee (ACUC) protocols MO23M346 (mice), AV23M340 (zebra finches), and AM23M342 (frogs). We purchased C57BL/6J mice from the Jackson Laboratory and used male offspring for experiments. We purchased adult male zebra finches from MD Exotic Birds and adult female Xenopus tropicalis from Xenopus I, and used the purchased animals for experiments. Tissue samples from *Octopus bimaculoides* were supplied by Cristopher Niell at the University of Oregon.

#### SNAIL and TRI probe design and preparation

SNAIL and TRI probes were designed according to previous STARmap and RIBOmap publications, with modifications made to incorporate the hybridization site for an Illumina-based sequencing primer.^8,10^ For genes with multiple isoforms, we identified regions conserved across all annotated isoforms and designed probes targeting these shared sequences. We selected unique target sequences using PICKY v2.2 to minimize off-target hybridization, and designed the hybridization arms of each probe pair to be 40–46 nucleotides in length. Each SNAIL and TRI probe set corresponding to a unique gene carried a randomly generated gene-specific ID sequence encoded in the padlock probe. We designed gene ID sequences with a minimum Hamming distance > 2 between genes. We designed four probes for each gene. For the Gene and Barcode Module experiments associated with **Fig. 2**, custom oPool oligo pools were ordered through IDT.

For the Gene Module cerebellum experiment (**Fig. 3**), we ordered a custom oligonucleotide pool from Twist Biosciences comprising up to four SNAIL probes per gene, targeting 1,761 genes, including all known transcription factors, genes involved in cellular migration and adhesion, a curated set of neuronal marker genes, and additional targets (**Supplemental Table 1**). We amplified these SNAIL probes using a previously published rolling circle amplification method with modifications to the protocol to improve the final oligo yield.^32^ To increase RCA amplification yield, we added 0.16 mg/mL T4 gene 32 protein (NEB M0300L) to each RCA step. We also replaced the nicking enzyme and primer sites for Nt.BspQI with those necessary for Nt.Alw1 digestion and digested the final RCA amplified samples with 0.3 Units (U) of Nt.Alw1 (NEB R0627S) and 0.3U of Nb.Btsl (NEB R0707S). To increase the yield of fully digested oligo products, we increased the number of nicking digestions to three total reactions, denaturing the nicking enzymes, rehybridizing the nicking primers, and adding 0.3 U of fresh nicking enzymes for every round. After digestion, we removed denatured proteins by pelleting the samples and performed ethanol precipitation to purify the amplified oligos. We rehydrated the samples with RNase-free water and calculated their final concentrations. We then lyophilized the amplified product and resuspended the pool to 200 µM in RNase-free water.

All probe sequences can be found in **Supplemental Table 1**.

### Sample preparation

#### Non-barcoded Tissue Samples

We deeply anesthetized C57Bl/6J mice (12 weeks old) and adult Zebra finches with an overdose of isoflurane, and adult Xenopus tropicalis in 2g/L of MS222 pH 7.0 (Sigma Aldrich E10521), and decapitated the animals. We dissected brains, immediately embedded them in O.C.T. compound, flash-froze the samples in isopentane cooled on dry ice and stored them at −80 °C until cryosectioning.

#### Barcoded Tissue Samples

We anesthetized mice with isoflurane (2.5% for induction and 1.5% for maintenance). We injected 400nL barcoded Sindbis virus (HZ120 packaged with jk114 as described in^33^) into the primary motor cortex (AP:+1.0mm, ML:±1.6mm, DV:-1.0mm) of 8- to 10-week-old C57Bl6/J mice. For fresh tissue samples, we dissected the brain 24 hours after injection, punched out the injection site, immediately embedded the tissue into O.C.T, froze it on dry ice and stored them at −80 °C until cryosectioning.

For fixed tissue, we deeply anesthetized injected C57BL/6J mice (12 weeks old) with isoflurane and performed transcardial perfusions using 1× PBS followed by 4% paraformaldehyde (PFA) (Boster Biological Technology, AR1068). We dissected the brain and post-fixed it in 4% PFA at 4 °C overnight and transferred them into 30% sucrose (Millipore Sigma, S5016) until it sank. We then embedded the brain in OCT, froze it on dry ice, and then stored the samples at −80 °C until cryosectioning.

For both fresh and fixed samples, we cryosectioned the samples into 20 µm slices, melted them onto Superfrost Plus Microscope Slides (Fisher Scientific, 1255015), and then refrozen on a dry-ice-cooled metal block. We stored the slides at −80 °C until library synthesis.

### BARseq3 Gene and Barcode Module

#### BARseq3 Gene Module Library Synthesis

For fresh frozen samples, we removed the slides from −80 °C and immediately immersed them in freshly prepared 4% PFA (Electron Microscopy Sciences, 50980492) in 1× PBS at room temperature for 60 min. We washed samples in 1× PBS and installed HybriWell-FL chambers (22 mm x 22 mm x 0.25 mm; Grace Bio-Labs). We then sequentially dehydrated the samples in 70%, 85%, and 100% ethanol for 5 min each. We added fresh 100% ethanol to the chambers and incubated the slides at 4 °C for 1.5 hours. After incubation, we washed the samples 5 times with PBSTwB [0.5%(vol/vol) Tween-20 (Millipore Sigma, P9416) in 1× PBS] and rehydrated the samples with 1× PBSTxG [0.5%(vol/vol) Triton X-100, 100 mM Glycine, 0.2 U/µl Superasin RNase Inhibitor in 1x PBS] for 15 min at room temperature.

For fixed samples, we treated fixed tissue slices with pepsin [0.2%(wt/vol) pepsin (MilliporeSigma, P7012) in 0.1 M HCl] for 3 min at room temperature and then washed the sections in 1× PBS. We then installed HybriWell-FL chambers and processed the sections using the same dehydration, ethanol incubation, washing and rehydration steps described above for fresh-frozen samples.

SNAIL and TRI probe library synthesis steps were adapted from STARmap^10^ and RIBOmap^8^. We denatured probe libraries by heating them to 90 °C for 3 min immediately prior to hybridization. We then added the hybridization solution [SNAIL probes: 50 nM per oligo for **Fig. 2B** and 25 nM per oligo for **Fig. 2J**; TRI probes: padlock and primer at 25 nM per oligo; splint probe at 2.5 µM per oligo, 2x SSC (Sigma-Aldrich, S6639), 10%(vol/vol) formamide (Calbiochem OmniPur, 75-12-7), 10%(vol/vol) Triton X-100 (Sigma-Aldrich, 93443), 20 mM ribonucleoside vanadyl complex (New England Biolabs, S1402S), 0.1 mg/ml Salmon Sperm DNA (Invitrogen, AM9680), 0.2 U/µl Superasin RNase Inhibitor (Invitrogen, AM2696)] to the chambers and incubated the samples in hybridization solution at 40 °C for 16-20 hours. After hybridization, we washed samples twice with PBSTwB at room temperature, followed by one wash in PBSTwBR [4× SSC, 0.2 U/µl Superasin RNase Inhibitor in PBSTwB] at 37 °C for 20 min, and two additional washes in PBSTwB at room temperature.

We prepared the ligation solution [1× T4 DNA ligase buffer, 0.1 mg/ml recombinant albumin (New England Biolabs, B9200S), 0.2 U/µl Superasin RNase Inhibitor (Invitrogen, AM2696), 2 U/µl T4 DNA ligase (Thermo Fisher Scientific, EL0011)] and then incubated the samples in the ligation solution for 5 hours at room temperature and then washed the samples twice with PBSTwB.

We performed rolling circle amplification (RCA) in RCA solution [1× Phi29 buffer (MCLab, PP-200), 0.2 mg/ml BSA (New England Biolabs, B9200S), 0.4 mM dNTPs (Thermo Fisher Scientific, R0192), 250 µM aminoally-dUTP (Invitrogen, AM8439), 1 U/µl Phi29 polymerase (MCLab, PP-200), 0.2 U/µl Superasin RNase Inhibitor (Invitrogen, AM2696), 5% (vol/vol) Glycerol (Millipore Sigma,65516)] at 25 °C overnight within a hybridization chamber.

After RCA, we washed samples once with PBSTwB and crosslinked the products in crosslinking solution [50 nM BS(PEG)9 (Broadpharm, BP-21504; stock prepared in DMSO (Thermo Fisher Scientific, D12345)), 20 mM RVC (New England Biolabs, S1402S) in PBSTwB]. We incubated samples at room temperature for one hour and washed samples once with 1 M Tris-HCl (pH 8.0). We then incubated the samples in fresh 1 M Tris-HCl (pH 8.0) containing 0.2 U/µL Superasin RNase inhibitor for 30 min at room temperature, and washed them twice with PBSTwB.

#### BARseq3 Barcode Module Library Synthesis

The following steps follow on directly from the Gene Module Library Synthesis, and are modified from BARseq2^21^. We added the reverse transcription solution [1× QuantumScript™ Reverse Transcriptase buffer, 0.2 mg/ml BSA, 1 U/µl RiboLock RNase inhibitor, 0.5 mM dNTPs, 1 µM barcode LNA primer, 20 U/µl QuantumScript™ Reverse Transcriptase] and incubated samples at 37 °C for 7 hours.

After reverse transcription, we washed samples once with PBSTwB and crosslinked the cDNA in 50nM BS(PEG)9 in PBSTwB for 1 hour at room temperature. We washed the samples once with 1 M Tris-HCl (pH 8.0). We incubated samples in fresh 1M Tris-HCl (pH 8.0) for 30 min at room temperature, and washed samples twice with PBSTwB.

We then prepared gap-filling mix [1× Ampligase buffer, 50 mM KCl, 0.1 µM barcode padlock, 0.05 mM dNTPs, 5% (vol/vol) Glycerol, 20%(vol/vol) formamide, 0.5 U/µl Ampligase (Biosearch Technologies, A0102K), 1 U/µl RiboLock RNase inhibitor, 0.001 U/µl Phusion DNA polymerase (ThermoFisher, F530S), 0.4 U/µl RnaseH (QIAGEN, Y9220L)]. We incubated samples in gap-filling mix at 37 °C for 20 min and then at 45 °C for 45 min. Then, we washed samples twice with PBSTwB.

We performed the second RCA in RCA mix [1× Phi29 buffer, 0.2 mg/ml BSA, 0.25 mM dNTPs, 125 µM aminoally-dUTP, 1 U/µl Phi29 polymerase, 5%(vol/vol) Glycerol] at 25 °C overnight. We then washed the samples once with PBSTwB and crosslinked the products again in 50 nM BS(PEG)9 in PBSTwB for 1 h at room temperature. We washed samples once with 1 M Tris-HCl (pH 8.0), incubated samples in fresh 1 M Tris-HCl (pH 8.0) for 30 min at room temperature, and then washed samples twice with PBSTwB.

### BARseq3 transcriptomics-only assay in adult mouse cerebellum

#### Glass-bottom plate preparation

We coated Glass-bottom plates (MatTek, P24G-1.5-13-F) with a Bind-Silane coating solution [0.1%(vol/vol) Bind-Silane (Sigma-Aldrich, 440159-100ML), 80%(vol/vol) ethanol, 2%(vol/vol) glacial acetic acid (Sigma-Aldrich, A6283-100ML) in ultrapure water (Invitrogen, 10977-023)]. We left the coating solution within the wells until it evaporated. We then rinsed the wells once with 100% ethanol, added 1 mg/mL Poly-D-lysine (Sigma-Aldrich, A-003-E) per well, sealed the plates with Parafilm, and incubated them overnight at 4 °C. We washed wells three times with RNase-free water (Invitrogen, 10977-023) and air-dried them before cryosectioning.

#### Tissue sectioning and fixation

We cryosectioned samples at 16 µm, melted them directly onto the glass well surface, and refroze them. After cryosectioning, we air-dried the sections at room temperature until translucent (~5 min) and fixed them in 4% paraformaldehyde (EM-grade PFA; Electron Microscopy Sciences, 15710-S, diluted in 1× PBS; Thermo Fisher Scientific, Gibco 70011044) for 10 min at room temperature and then washed samples three times with 1× PBS. We then added ice-cold 100% methanol to the samples, replaced them with fresh ice-cold 100% methanol, and placed the samples at −80 °C until library synthesis.

#### BARseq3 Gene Module Transcriptomic-Only Library Synthesis

We removed the samples from −80 °C and allowed them to come to room temperature. We rehydrated samples in 1× PBSTwG [0.1%(vol/vol) Tween-20 (Millipore Sigma, P9416), 100 mM Glycine, 0.2 U/µl Superasin RNase Inhibitor, 0.1 mg/mL yeast tRNA (Invitrogen, AM7119) in 1× PBS] for 15 min at room temperature. We denatured SNAIL probe libraries by heating them at 90 °C for 3 min immediately prior to hybridization. We prepared the hybridization buffer as described above, substituting salmon sperm DNA for yeast tRNA and using 1.5 nM per SNAIL oligo. We then incubated the samples in hybridization buffer, covered the well plate with parafilm, and placed it in a hybridization chamber, incubating at 40 °C for 40 hours with gentle agitation. After hybridization, we washed the samples with PBSTwR [0.1%(vol/vol) Tween-20 in 1x PBS with 0.2 U/µl Superasin RNase Inhibitor] at room temperature twice for 20 minutes at room temperature, and then treated with 4X SSC in PBSTwR (4x SSC, 0.2 U/µl Superasin RNase Inhibitor in PBSTwG) for 20 minutes at 37°C. We performed ligation as previously stated, with an incubation time of 2 hours at room temperature, and then we washed the samples twice with PBSTwR for 20 mins each. We performed RCA in RCA solution (1× Phi29 buffer, 0.2 mg/ml BSA, 0.25 mM dNTPs, 125 µM aminoally-dUTP, 1 U/µl Phi29 polymerase, 0.2 U/µl Superasin RNase Inhibitor) for 30 min at 4 °C, followed by a 2-hour incubation at 30 °C. We crosslinked the rolonies as described above without the Superasin RNase Inhibitor.

### BARseq3 Sequencing Module

#### Sequencing primer hybridization

We prepared the stripping buffer [40%(vol/vol) formamide (Calbiochem OmniPur, 75-12-7),1%(vol/vol) Triton X-100 (Sigma-Aldrich, 93443)]. We incubated the samples in stripping buffer on a 60 °C heated metal block for 5 min, removed the samples to cool to room temperature, and repeated the incubation once more. We washed samples twice with PBSTwS [2%(vol/vol) Tween-20 in 1× PBS]. We then prepared the sequencing primer hybridization solution [10%(vol/vol) formamide, 2x SSC] and washed the samples once with the sequencing primer hybridization solution. We then incubated the library in a sequencing primer hybridization solution with 1 uM sequencing primer for 10 min at room temperature. All sequencing primer sequences can be found in **Supplemental Table 1**. After hybridization, we washed the samples once in a sequencing primer hybridization solution, then washed twice in PBSTwS.

#### Sequencing cycle 1

All Illumina sequencing reagents [Incorporation buffer, Incorporation Reaction Mixture (IRM), Cleavage reaction mix (CRM)] were taken from an MiSeq Reagent Nano 500 cycle Kit v2 (Illumina, MS-103-1003). For all sequencing washes performed at 60 °C on a heated metal block, we cooled samples to room temperature before exchanging solutions.

We incubated samples twice in incorporation buffer at 60 °C for 3 min each, washed samples once in PBSTwS at room temperature, incubated samples in Iodoacetamide blocker solution [2.6mg/mL Idoacetamide (Thermo Fisher Scientific, A39271) in PBSTwS] at 60 °C for 3 min, and rinsed samples again in PBSTwS. We washed samples twice in incorporation buffer, followed by two incubations in IRM at 60 °C for 3 min each. We then washed the samples four times with PBSTwS at 60 °C for 3 min each. We incubated samples with 1:100 Blue Nissl (Invitrogen, N21479) in PBSTwB for 20 mins and washed 3 times with PBSTwB before imaging.

#### Sequencing cycles 2-n

We washed samples twice in the incorporation buffer. Then, we incubated samples twice in CRM at 60 °C for 3 min each. We washed samples twice with incorporation buffer, rinsed samples in PBSTwS at room temperature, incubated samples in Iodoacetamide blocker solution [2.6 mg/mL Idoacetamide (Thermo Fisher Scientific, A39271) in PBSTwS] at 60 °C for 3 min, and rinsed samples again in PBSTwS. We then washed samples twice in the incorporation buffer at room temperature, followed by two incubations in IRM at 60 °C for 3 min each. Finally, we washed samples four times in PBSTwS at 60 °C for 3 min each before imaging. In the experiments for **Fig. 2**, we performed a total of five Gene Module sequencing cycles, except for the *Pcp2*/*Malat1* experiments, where we sequenced only two cycles.

After Gene Module sequencing, we stripped the gene sequencing primer using a higher formamide stripping solution [(80%(vol/vol) formamide (Calbiochem OmniPur, 75-12-7),1%(vol/vol), Triton X-100 (Sigma-Aldrich, 93443), 2x SSC]. We then hybridized the barcode sequencing primer and performed 10 additional sequencing cycles as described above.

For the BARseq3 Gene Module only data (**Fig. 3**), 2.6 mg/mL Iodoacetamide was replaced with 0.4% MMTS (Thermo Fisher Scientific, 23011) diluted in PBSTx (0.1% Triton X-100 in 1× PBS), and we used PBSTx in place of PBSTwS for all sequencing cycles. After the first sequencing cycle, we incubated the samples in 1:1000 DAPI (Thermo fisher, 62248) in PBSTx for 5 min and then incubated samples with 1:100 Blue Nissl (Invitrogen, N21479) in PBSTx for 20 min. We performed 10 sequencing cycles for this data.

### BARseq2 experiments

We designed BARseq2 padlock probes using the Padlock designer code used in BARseq2^21^ and synthesized 12 padlocks per gene as oPool oligo pools from IDT. We performed all the BARseq2 experiments according to the published BARseq2 protocol (https://dx.doi.org/10.17504/protocols.io.81wgbp4j3vpk/v2). All probe sequences can be found in **Supplemental Table 1**.

### Immunostaining

After completion of the 10th sequencing cycle for Gene Module only data, we performed an 11th sequencing cycle. We then washed samples in 1× PBS and performed the immunostaining as follows. We treated sections with PBSTx for 10 min at room temperature, then blocked in blocking buffer [5%(vol/vol) normal goat serum in PBSTx] for 1 hr at room temperature. We incubated the samples with ZsGreen1 antibody (Takara, 632474; 1:500 in blocking buffer) overnight at 4 °C in a humidified chamber. We washed the samples three times in PBSTx (5– 10 min each) and incubated samples with Alexa Fluor™ 488–conjugated secondary antibody (Thermofisher Scientific, A32731TR; 1:1000 in blocking buffer) for 1 hour at room temperature protected from light. We then washed sections three times for 5–10 min each in PBSTx, rinsed once in 1× PBS, and imaged.

### Microscope and Imaging Specifications

We imaged samples using a Crest X-Light V3 spinning-disk confocal mounted on a Nikon Ti2-E using a 25x silicone-oil (NA 1.05; **Fig. 2**) or 40x water-immersion objective (NA 1.25; **Fig. 3**). Illumina fluorophores were excited at 514 (G/T) and 640 nm (A/C). Simultaneous two-channel acquisition used two Hamamatsu ORCA-FusionBT cameras synchronized via hardware TTL triggering from the microscope controller using NIS-Elements software. Fluorescence separation was achieved using a ZT405/514/635rpc dichroic with a ZET532/640m emission filter, and camera cube containing a T637/105dcbp-UT2 camera dichroic and a ET550/40+752/95m cleanup filter.

### Computational analysis pipeline - image processing, decoding, and assignment

We implemented a custom computational pipeline to process four-channel, multi-cycle sequencing images, detect and decode amplicons, and assign gene identities to each cell. We first performed maximum projection of the image stacks along the z axis. We aligned imaging channels using a homography transform and then registered all sequencing cycles to the cycle containing the DAPI or Nissl signal by using an affine transform for coarse alignment followed by Demons^34^ for nonlinear refinement. We then performed spectral unmixing, min– max normalization and rolling-ball background subtraction (radius = 15 for **Fig. 2** analyses; radius = 10 for **Fig. 3** analyses). For transcriptomic reads, we detected fluorescent amplicons using Bardensr^24^, which identifies spots by convolving images with a point-spread function and assigns confidence scores to candidate gene IDs based on matches to the gene codebook. We used a confidence threshold of 0.7 for **Fig. 2** analyses and set the false discovery rate (FDR) to 10% for **Fig. 3** analyses. The FDR was generated using a codebook augmented with synthetic “false gene IDs” that do not occur in any probes but have a Hamming distance of two to all other gene IDs. We performed cell segmentation based on DAPI or Nissl images using Cellpose^35^ and assigned gene calls to cells based on spatial overlap with the corresponding cell masks to generate the expression matrix. To process BARseq3 data in **Fig. 2**, we aligned the Nissl signal masks between the Barcode Module and the Gene Module and applied the transformation to barcode images. We then segmented barcoded cells in the barcode images using Cellpose^35^ and excluded cells with a barcode mask area < 1,000 pixels (~68µm^2^). For base calling, we assigned the nucleotide at each sequencing cycle based on the maximum intensity nucleic-acid fluorescence signal within each cell mask. Finally, we mapped the spatial coordinates of barcoded cells onto the Gene Module cell spatial location to link each barcoded cell to its corresponding gene-expression profile.

### Gene signal specificity experiments and analysis

We designed probes targeting *Pcp2* (Purkinje cell-specific) and *Malat1* (nuclear-specific) and performed BARseq3 Gene Module-only experiments on fresh-frozen mouse brain tissue using probe concentrations of 5, 10, and 20 nM on glass slides and 25x imaging as described above. We performed parallel experiments targeting *Pcp2* and *Malat1* using the standard BARseq2 protocol^21^. The two gene IDs were chosen such that they were assigned to the C channel and G channels in the first round, respectively, to minimize interference from color bleeding, and were assigned to G and A in the second round. For all analyses, we performed two biological replicates with two sections per replicate and selected two FOVs per section from the cerebellar cortex (*Pcp2, Malat1* experiments) and brainstem (*Pcp2* experiments).

For *Malat1* analysis, we selected sparse cerebellar regions of interest (4 sections, 2 FOVs each) in the molecular layer from each image using DAPI-guided region selection. We locally aligned G channel images, which contained *Malat1* signal, to the corresponding DAPI/Nissl reference images using small rigid transformations. We segmented nuclear masks from DAPI images using Cellpose^35^. We then defined cell territories from Nissl images using nucleus-seeded segmentation and expanded territories until they reached neighboring cells. For each cell, we calculated a *Malat1* signal-to-noise ratio as the mean *Malat1* intensity inside the expanded DAPI mask divided by the mean *Malat1* intensity in the surrounding non-nuclear territory within the expanded Nissl mask. We compared conditions using Welch’s t-tests with Bonferroni correction.

For *Pcp2* spot-based cerebellar analysis, we segmented Purkinje cells from cerebellar FOVs. We identified candidate Purkinje cells from C channel spot structure, which contained only *Pcp2* signal, compared these with Nissl segmentation, and generated Purkinje-cell masks. We counted *Pcp2* puncta using skimage.peak_local_max() in min-max normalized images and retained only bright *Pcp2* spots with background-subtracted intensity greater than 300. For each biological slice and replicate, we calculated the average number of *Pcp2* spots per Purkinje cell across all available cerebellar FOVs. We plotted these slice-by-replicate measurements as four datapoints per condition (slice1 rep1, slice1 rep2, slice2 rep1, slice2 rep2) and compared conditions using Welch’s t-tests with Bonferroni correction. We repeated this analysis for brainstem FOVs, finding random regions that are matched in size to Purkinje cells and performing the same slice-level measurements.

### BARseq3 transcriptomics-only in adult mouse cerebellum analysis

#### Cell Type Analysis

We generated cell-by-gene count matrices with associated spatial metadata using the computational pipeline described above. We removed counts fewer than 3 or greater than 400 prior to analysis. We log-normalized and scaled gene counts for each cell and then performed principal component analysis (PCA). We retained the first 30 principal components for downstream analysis. We constructed a neighborhood graph using 12 nearest neighbors and embedded cells using UMAP with a minimum distance parameter of 0.1. We performed clustering with the Leiden algorithm at a resolution of 1.3 to identify major cell populations. We then subsetted excitatory and inhibitory neurons and re-clustered the cerebellar nuclei population separately at a resolution of 0.2 to resolve excitatory vs inhibitory neuron subtypes. Finally, we assigned cell type identities based on the expression of established marker genes.

#### Replicate Analysis

To assess reproducibility, we processed serial sections of adult mouse cerebellum independently using the BARseq3 transcriptomics-only assay. We quantified gene counts in each section and compared gene expression profiles across sections. This analysis revealed a high correlation in gene counts between replicate sections.

#### Signal-to-Noise Ratio Analysis

We quantified signal-to-noise ratios across sequencing cycles using manual pixel selection from ten representative FOVs. For each FOV and imaging cycle, ‘signal’ pixels were manually selected as the brightest pixel from clearly identifiable amplicons, and background pixels were selected from nearby regions lacking visible amplicon signal. We extracted raw fluorescence intensities directly from the images. For each FOV and cycle, we calculated SNR as the difference between the signal intensity and background intensity divided by the standard deviation of the background intensity. Mean SNR was calculated across FOVs within each sequencing cycle across the ten FOVs. We used the same methods to calculate SNR for **Fig. 2D** and **Sup. Fig. 1B-E** from 2-3 FOVs.

## DATA AND CODE AVAILABILITY

Analysis code is available on GitHub (https://github.com/kebschulllab/BARseq3). Raw and processed data files are available on Zenodo (10.5281/zenodo.19024678).

## ACKNOWLEDGMENTS

We thank Cristopher Niell and Angelique Allen for supplying octopus tissue samples. We also appreciate valuable feedback on this manuscript by Jean Fan, Xiaoyin Chen, as well as Fan and Kebschull lab members. This work was supported by NIH grants DP1DA056668, R34132027, RF1AG078378, U01NS132161, R01DA054374, and R01DA056599 to JMK; the Packard, Klingenstein Simons, and Sloan Fellowships to J.M.K, and a Kavli Distinguished PhD Fellowship to M.M.G.A., an NSF GRFP to C.S., and NIH Training grant T32DC000023 to C.D.W.

## AUTHOR CONTRIBUTIONS

H.Q., M.M.G.A, D.Z.F-G., and J.M.K. conceptualized the study. H.Q. and L.W. performed the experiments to integrate Gene and Barcode Modules. M.M.G.A. and D.Z.F-G. performed the experiments to integrate Gene and Sequencing Modules with support from I.D.R.. H.Q. performed the translatomics experiments. M.M.G.A. performed the cerebellar experiments. M.M.G.A. led the development of the computational pipeline to analyze BARseq3 datasets, with help from R.H., C.D.W., and D.Z.F-G.. C.S. performed BARseq3 experiments in fixed tissue. H.Q., M.M.G.A, and D.Z.F-G. performed all other experiments. M.M.G.A drafted the initial manuscript. H.Q., M.M.G.A, D.Z.F-G., and J.M.K. prepared the final version.

## AI STATEMENT

AI tools were used for language editing and proofreading during manuscript preparation. All scientific content, analyses, interpretations, and conclusions were generated and verified by the authors.

## SUPPLEMENTAL FIGURES

**Supplemental Figure 1:**
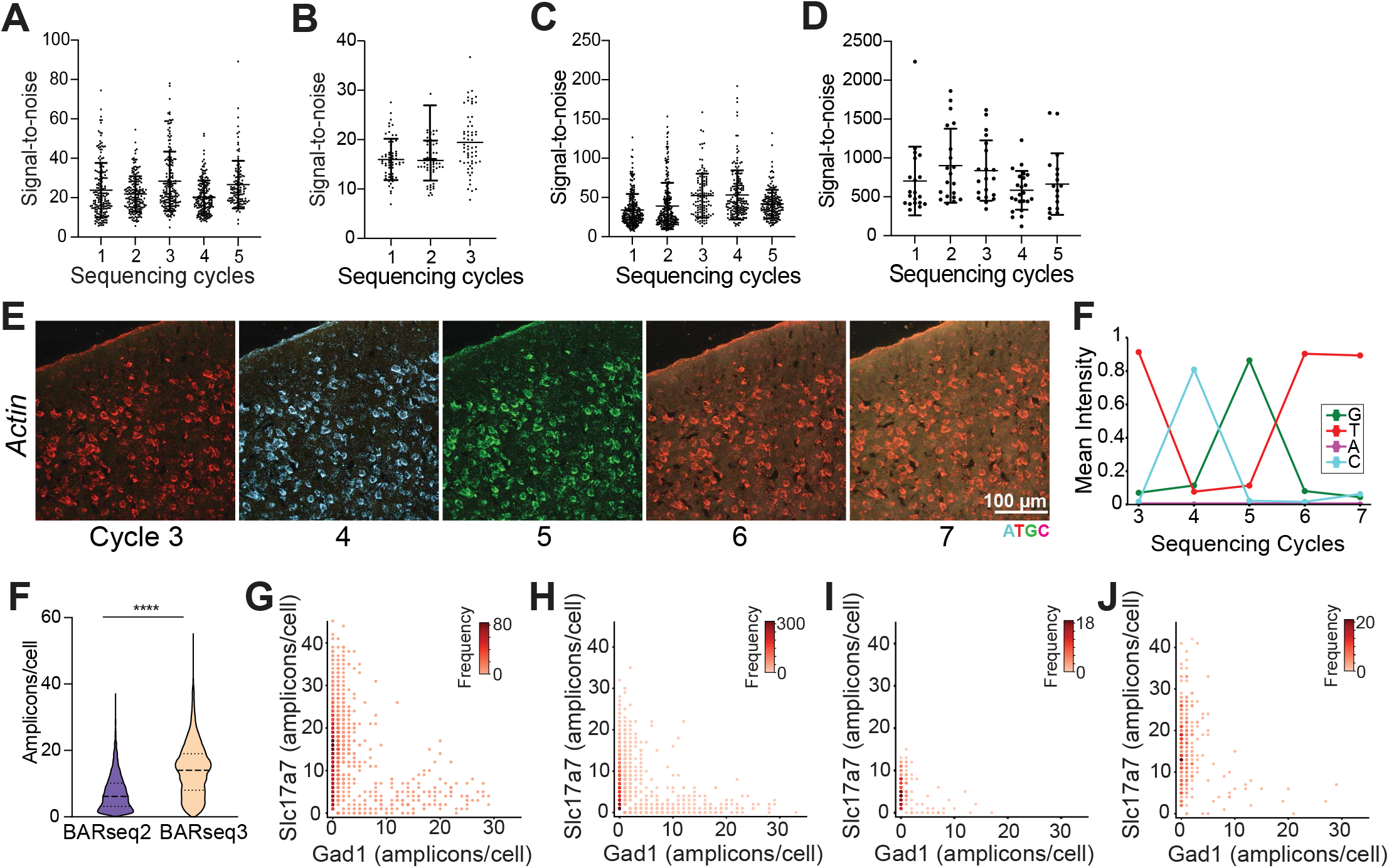
BARseq3 produces robust, sequence-able amplicons allowing for accurate gene decoding. **(A, B)** Signal-to-noise of BARseq3 transcriptome **(A)** and translatome **(B)** amplicons remains stable across sequencing cycles. Absolute value of signal-to-noise is largely determined by the imaging setup (here, 25X silicone oil objective imaged through a HybriWell-FL chamber; compare **Fig. 3**, with more optimal imaging conditions). **(C,D)** Signal-to-noise of BARseq2 transcriptome amplicons **(C)** and barcodes **(D)** remains stable across sequencing cycles. **(E,F)** Gene Module amplicons in an experiment probing for a single gene (β-Actin) change fluorescence across sequencing cycles **(E)** with low levels of sequencing lag **(F)**. Quantification of *Slc17a7* and *Gad1* gene amplicons in non-barcoded cells shows that BARseq3 produces more amplicons than BARseq2. (BARseq2 n=3140 cells, BARseq3 n=2322, Student’s t-test, ****pvalue < 0.0001). **(G-J)** *Slc17a7* and *Gad1* amplicons in non-barcoded BARseq3 **(G)** and BARseq2 **(H)**, as well as barcoded BARseq2 **(I)** and BARseq3 cells **(J)**, are localized to different cells.

**Supplemental Figure 2:**
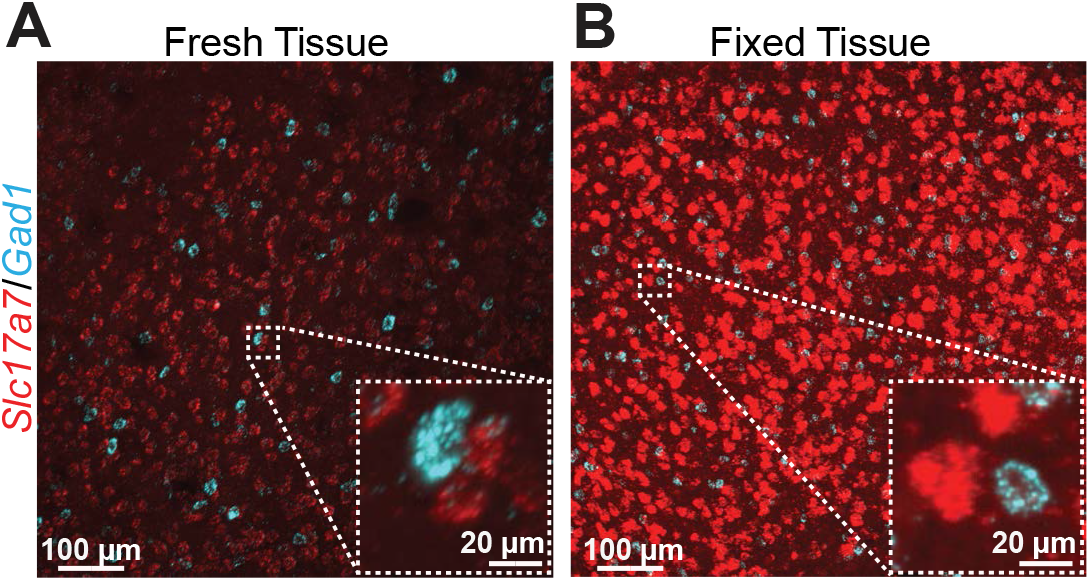
BARseq3 in fresh and fixed tissue. **(A)** Fresh tissue samples within the mouse cortex using BARseq3. **(B)** Pepsin treatment of PFA fixed tissue recovers Gene Module amplicons during library synthesis.

**Supplemental Figure 3:**
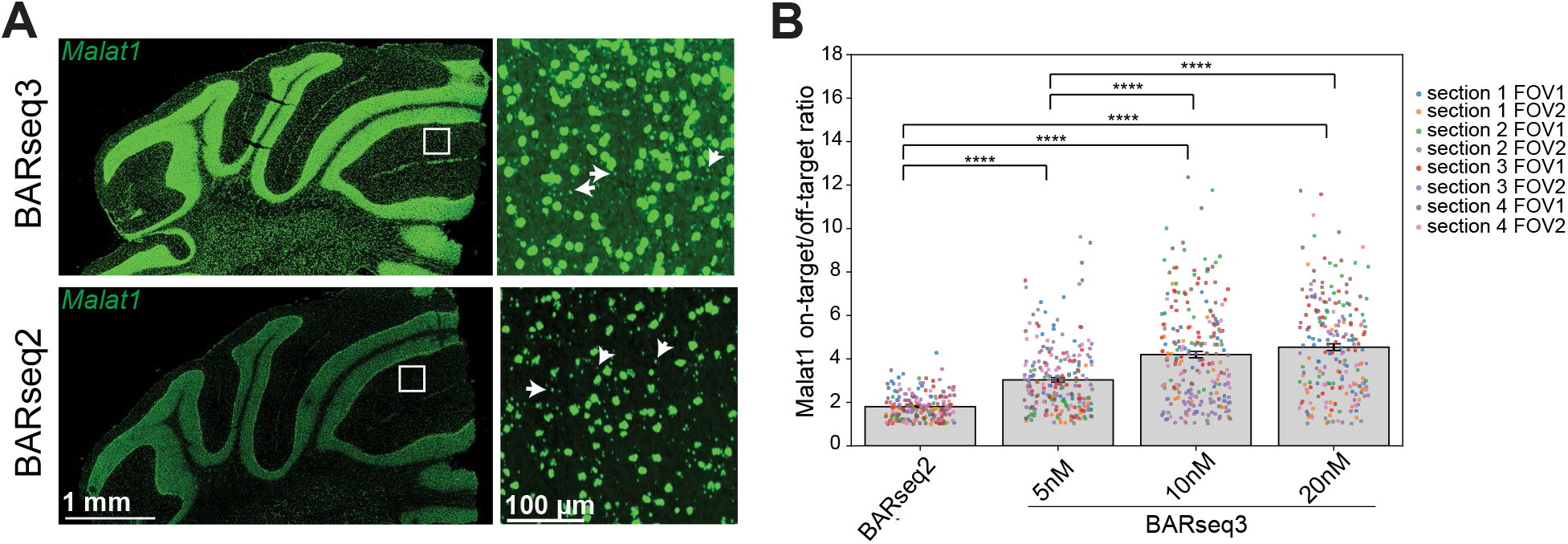
Specificity of Malat1 expression in BARseq3. **(A)** Representative images of the *Malat1* specificity experiment (10 nM, BARseq3), arrows highlight off-target amplicons outside the nucleus. **(B)** BARseq3 *Malat1* on-target signal/off-target signal is higher than BARseq2 and increases as probe concentration increases (n = 8, FOVs each, Welch’s t-test with Bonferroni correction, ****pvalue < 1.67e-5).

**Supplemental Figure 4:**
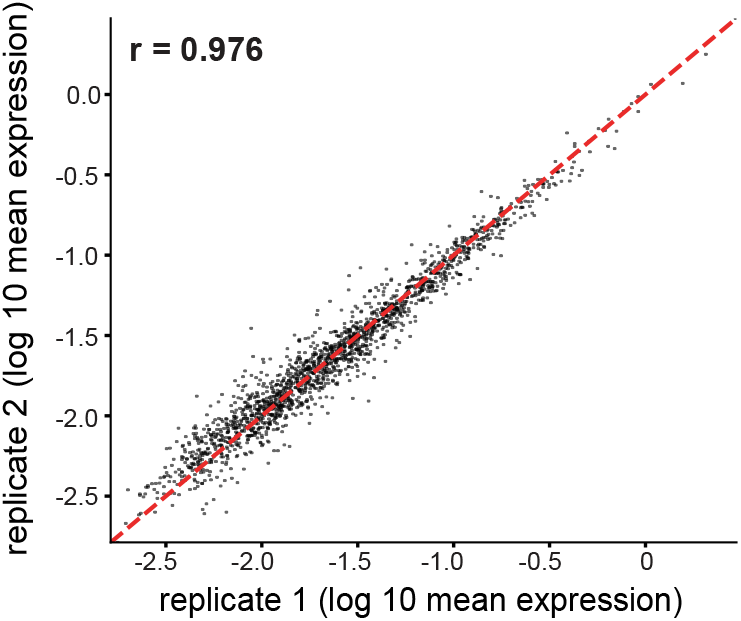
Adjacent serial sections show high gene expression reproducibility. Scatterplot of log 10 gene expression per gene for serial section 1 (replicate1, x-axis) and serial section 2 (replicate 2, y-axis). The dashed red line indicates equal values between both replicates.

**Supplemental Figure 5:**
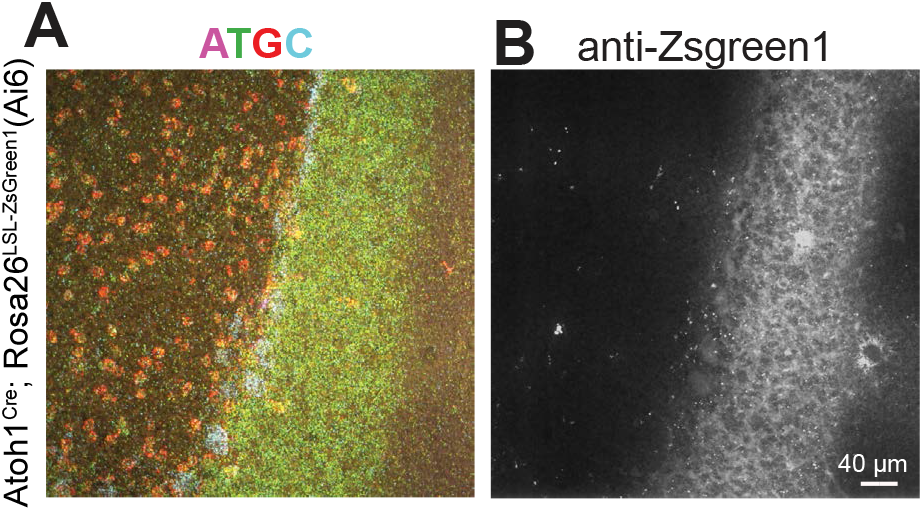
BARseq3 is compatible with immunostaining. **(A,B)** anti-Zsgreen1 antibody staining of genetically labeled cells in the Atoh1-lineage after the last cycle of the gene library Sequencing Module in a mouse cerebellar section **(A)** produces strong signal at in the expected cell population (granule cells, **B**).

